# VipA N-terminal linker and VipB-VipB interaction modulate the contraction of Type VI secretion system sheath

**DOI:** 10.1101/152785

**Authors:** Maximilian Brackmann, Jing Wang, Marek Basler

## Abstract

Secretion systems are essential for bacteria to survive and manipulate their environment. The bacterial Type VI Secretion System (T6SS) generates the force needed for protein translocation by the contraction of a long polymer called sheath, which is composed of interconnected VipA/VipB subunits forming a six-start helix. The mechanism of T6SS sheath contraction and the structure of its extended state are unknown. Here we show that elongating the N-terminal VipA linker or eliminating charge of a specific VipB residue abolished sheath contraction and delivery of effectors into target cells. The assembly of the non-contractile sheaths was dependent on the baseplate component TssE and mass-spectrometry analysis identified Hcp, VgrG and other components of the T6SS baseplate specifically associated with stable non-contractile sheaths. The ability to lock T6SS in the pre-firing state opens new possibilities for understanding its mode of action.

## Introduction

Various protein nanomachines have evolved to translocate macromolecules across biological membranes. A subset of these nanomachines is composed of a rigid tube surrounded by a contractile sheath attached to a baseplate. The sheath is initially assembled in a high energy, extended state and then quickly transits to a low energy, contracted state, which results in physical puncturing of the target membrane associated with the baseplate. The bacterial Type VI secretion system (T6SS) uses this mechanism to deliver proteins across membranes [1–5].

Assembly of all contractile nanomachines starts by formation of a baseplate, which initiates polymerization of the inner tube. The tube serves as a template for quick assembly of the extended sheath [6–10]. Unlike other contractile systems, the assembly of T6SS baseplate requires the interaction with a membrane complex, which anchors the system to the cell envelope [11,12]. Two different classes of TssA molecules are important for initiation of T6SS sheath assembly and its elongation [13,14], which progresses across the whole cell and thus allows the use of live-cell fluorescence microscopy to monitor the sheath dynamics [1,12,15–17].

The sheath can be described as a six-start helix or as a stack of rings composed of six subunits, which are interconnected by N- and C-terminal linkers in the inner layer of the sheath. Inner layers of sheaths of contractile phage, R-type pyocin and T6SS likely maintain the connectivity of the sheath during contraction and are evolutionarily related to each other. However, the surface exposed domains are distinct [9,18–23]. Specifically, T6SS sheath contains a specific surface exposed Domain 3 of unknown structure [20,22,23] but plays a crucial role in sheath recycling.

The mechanism of T6SS sheath contraction is unknown because the sheath contracts during isolation from cells and thus the extended sheath is yet to be analyzed in detail [1].

Here, we identified two structural features of the T6SS sheath that play a critical role in its contraction. Using live-cell fluorescence microscopy, we show that a single negatively charged residue located on the surface of the middle domain of the T6SS sheath is critical for sheath contraction but not sheath assembly. We further show that VipA N-terminal linker structure is critical for sheath contraction. Insertion of two and more amino acid residues into this linker completely abrogated contraction and allowed us to isolate non-contractile sheaths from cells for mass-spectrometry and electron microscopy analysis. This analysis revealed that non-contracted sheaths are stably associated with Hcp tube and components of T6SS baseplate. Overall, our analysis shows that conserved structural features are involved in sheath contraction and provides insights into T6SS assembly and mode of action.

## Results

### VipB residue D333 is important for contraction *in-vivo*

Interactions of charged residues were previously suggested to be important for contraction of T4 phage sheath and R-type pyocin sheath [18,21]. Analysis of interfaces that are expected to be present only in the contracted form of the T6SS sheath of *V. cholerae* suggested that VipB residues K223 and D333 located on two different VipB-VipB interfaces, one between protomers of a single sheath strand and the second between protomers on two adjacent strands, significantly contribute to the stability of the contracted structure [22]. This suggests that energy released by forming these interactions could be contributing to driving the sheath contraction.

To test this hypothesis, we mutated K223 and D333 to alanine and expressed the mutated *vipB* in a *vipA-msfGFP* background to allow for monitoring of sheath dynamics in live-cells. K223A mutation impaired assembly of the sheath and no elongated sheath were observed (Fig. 1b). This was also reflected in the inability of the VipB-K223A mutant to kill target cells (Fig. 1a). Interestingly, VipB-D333A mutant assembled into long non-dynamic sheath structures that were stable *in-vivo* over more than one hour of imaging (Fig. 1c and Supplementary Video S1 and S2). The number of sheath structures per cell was comparable to the number of structures assembled in wild-type cells suggesting that D333 residue is not critical for sheath assembly. Importantly, VipB-D333A mutation completely blocked target cell killing (Fig. 1a).

**Figure 1.**
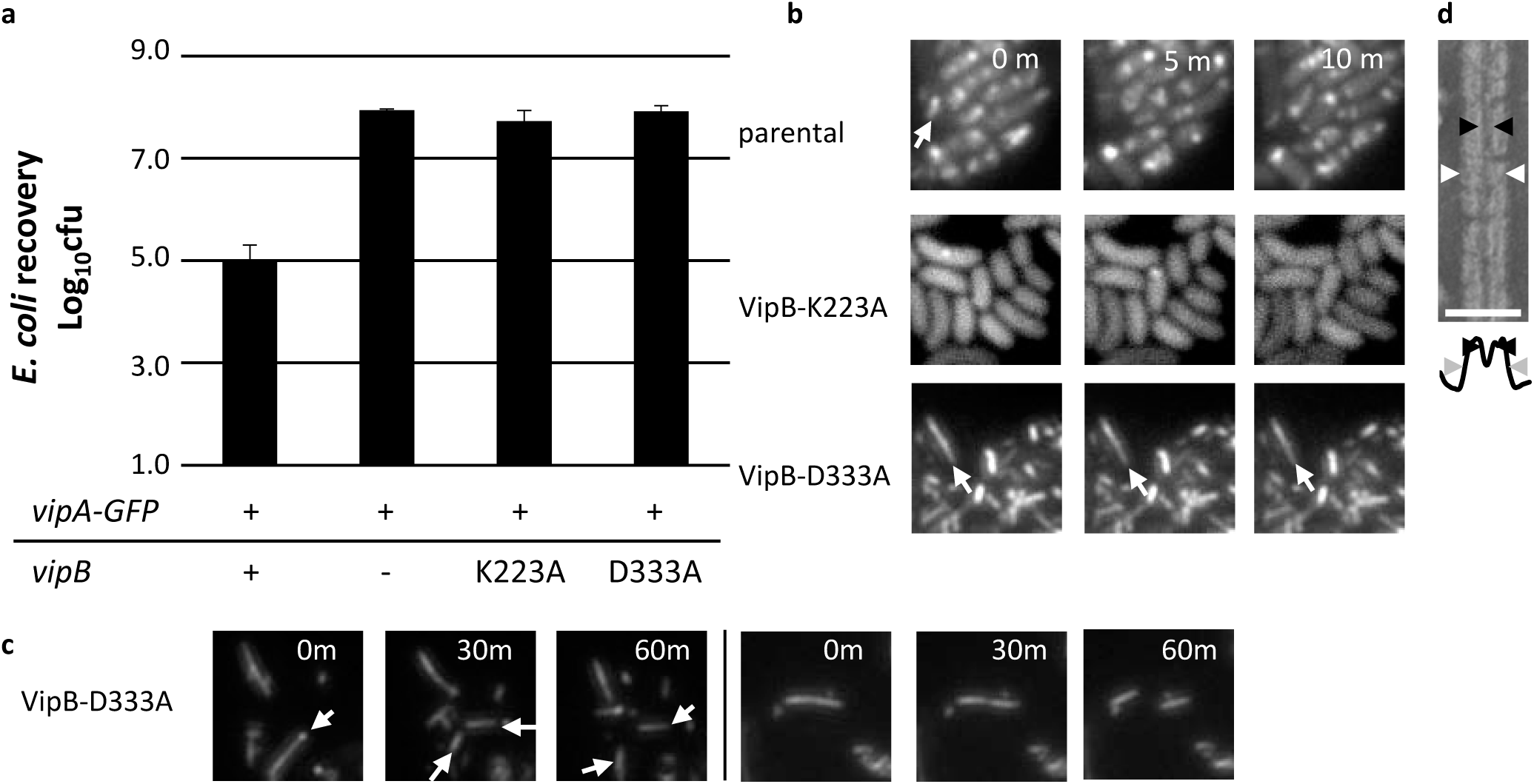
Charged residues of VipB are essential for sheath assembly and contraction a. *E. coli* survival (± SD, N=3) after 3h competition with indicated *V. cholerae* strains in a 1:10 ratio on plate. **b** Fluorescence timelapse images of *V. cholerae vipA-msfGFP, vipB^-^* complemented with *vipB*-K223A, *vipB*-D333A on pBAD24. **c** Long timelapses of *V. cholerae vipA-msfGFP, vipB^-^* complemented with *vipB-*D333A on pBAD24. **d** Electron micrograph of purified VipB-D333A sheath and below a plot of the summed intensities. The inner diameter is marked with black arrowheads and the outer diameter with grey arrowheads.

### VipA linker is critical for sheath contraction *in-vivo*

Recent atomic models of T6SS sheaths in a contracted state and structures of the R-type pyocin in an extended and contracted state identified intermolecular linkers important for sheath function [20–22]. Interestingly, the N-terminal linker of the R-type pyocin sheath is more stretched in the contracted sheath than in the extended sheath (Supplementary Fig. 1). T4 phage sheath was suggested to contract from the baseplate in a wave of sequentially contracting rings [24]. We hypothesized that stretching of the N-terminal sheath linker upon contraction suggests that the contraction of a basal ring of sheath results in pulling on the VipA N-terminal linker of the next ring, which in turn triggers its contraction. Such a mechanism would lead to propagation of contraction through the whole sheath. We decided to test this hypothesis by generating a series of mutant *V. cholerae* T6SS sheaths with longer VipA linkers that potentially hinder propagation of contraction. We inserted 1-7 amino acids of the native “AEVELPL” sequence of the linker after the residue 25 of VipA wild-type protein (labeled here as VipA-N1 to VipA-N7). To monitor the assembly and contraction, we fused the VipA and its variants to msfGFP and expressed it from pBAD24 plasmid in the absence of chromosomal *vipA* [1,22].

All mutant T6SS sheaths assembled with a frequency similar to the wild-type sheath (Fig. 2b), however, frequency of sheath contraction was strongly dependent on the linker length (Fig. 2b, Supplementary Video S3). Whereas insertion of one amino acid (VipA-N1) had almost no effect on sheath dynamics, an elongation by two or more amino acids (VipA-N2-7) reduced the fraction of sheaths that contract during 5 minutes from 50% (of 159 structures counted, 85 contracted) to 0% (of 204 structures counted, none contracted). Many of these mutant sheaths were stable over one hour of imaging (Fig. 2c and Supplementary Video S4). Sheaths that occasionally broke after extensive bending caused by movement and growth of cells were however quickly disassembled (Supplementary Video S4). VipA with “AGAGA” sequence inserted, labeled as VipAN5(GA), also assembled stable full-length sheaths (Fig. 2b, Supplementary Video S3), suggesting that VipA linker length, but not its sequence, is specifically critical for sheath stability. Furthermore, the killing of *E. coli* MG1655 by *V. cholerae* T6SS (Fig. 2a) was strongly dependent on the length of the VipA linker. Whereas an extension of this linker by one amino acid had no effect on the killing efficiency, an elongation by two amino acids completely abolished the killing of target cells.

**Figure 2.**
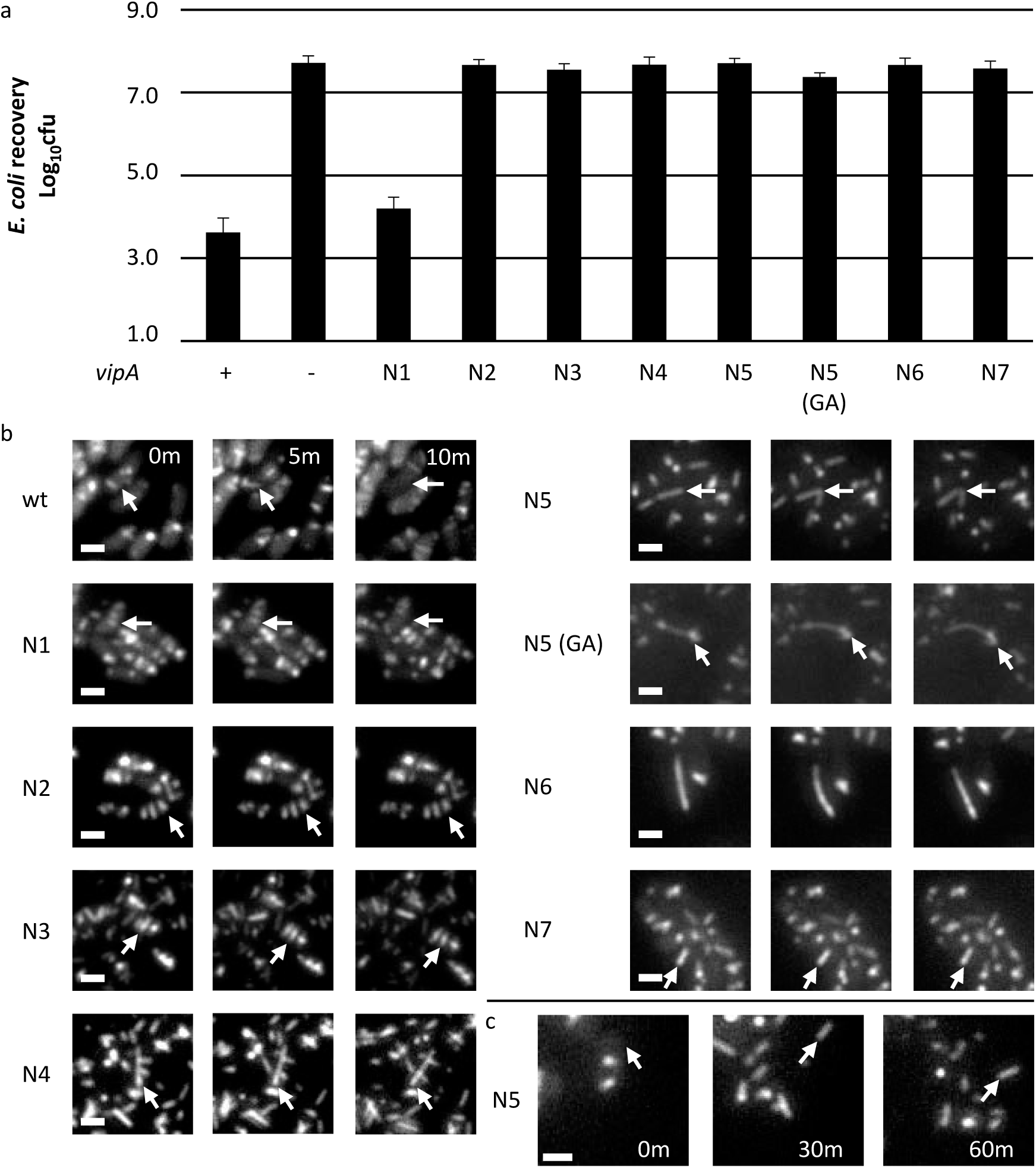
Length of N-terminal linker of VipA controls sheath contraction. **a** *E. coli* survival (± SD, N=3) after 3h competition with indicated *V. cholerae* strains in a 1:10 ratio on plate. **b** Fluorescence timelapse images of *V. cholerae vipA^-^* complemented with indicated msfGFPtagged-*vipA* variants on pBAD24. **c** Long timelapse of *V. cholerae vipA^-^* complemented with *vipA-N5-msfGFP* on pBAD24. Scale bars are 1 μm.

### Stable sheaths assemble from baseplate and around Hcp

To analyze the stable mutant sheaths in more detail, we isolated them by using an approach similar to the one used for the isolation of the wild-type contracted sheath [1,22]. Mutant sheaths were expressed in non-flagellated *V. cholerae* strain, cells were lysed and sheaths were purified from soluble proteins and cell debris using ultra-centrifugation. The isolated sheaths were analyzed by negative staining electron microscopy.

Analysis of VipB-D333A sheath sample revealed partially fragmented hollow structures with an outer diameter of 260 nm, thus resembling contracted sheaths (Fig. 1d). This suggests that during isolation the VipB-D333A sheaths contract and the D333A mutation destabilizes the contracted structure and this leads to partial fragmentation. Similarly, VipA-N2 sheaths closely resembled the wild-type contracted sheaths as they appeared hollow, had the inner diameter of 100 Å and the outer diameter of 260 Å (Fig. 3a, b). Interestingly, VipA-N3, VipA-N5 and VipA-N5(GA) mutant sheath diameters were ≈200 Å and thus narrower than wild-type (Fig. 3a, b). Importantly, uranyl acetate stain was clearly unable to penetrate the sheaths, suggesting that VipA-N3, VipA-N5 and VipA-N5(GA) mutant sheaths were filled with additional protein (Fig. 3a, b).

**Figure 3.**
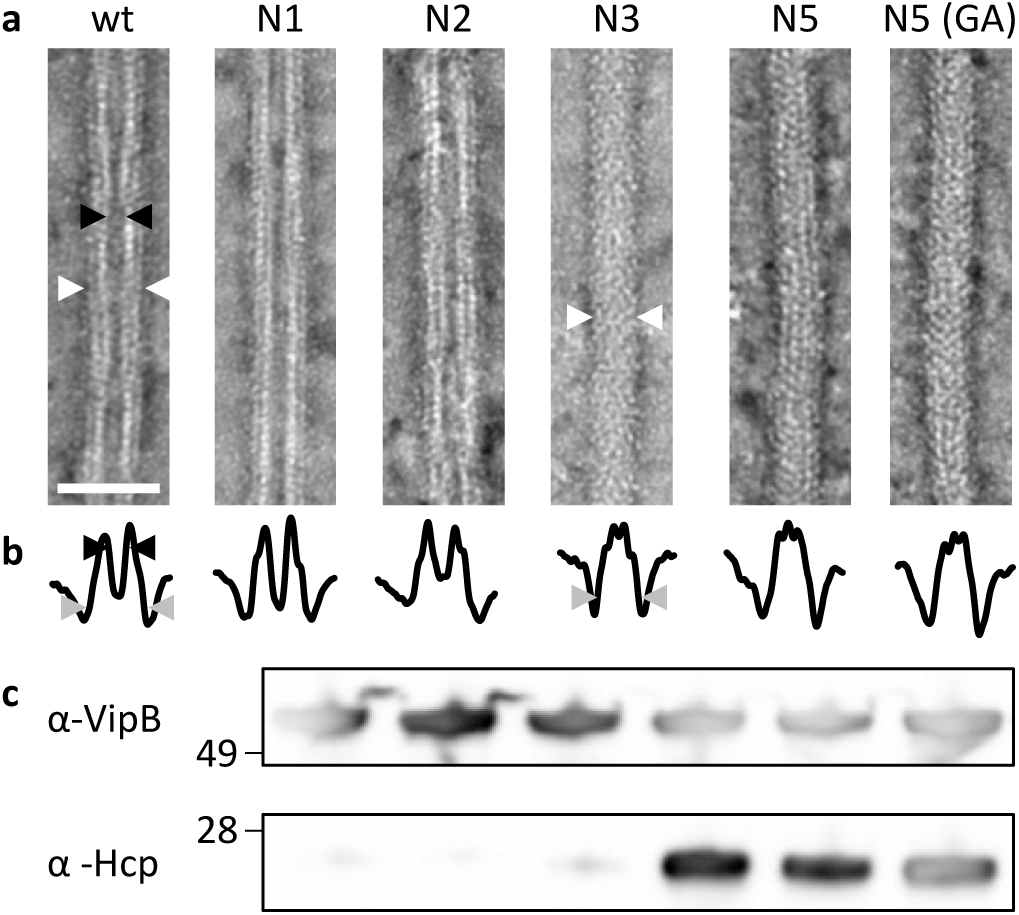
Hcp is enriched in VipA mutants with a linker elongated by three or more amino acids. **a** Electron micrographs of sheath samples. Wild-type sheath and those with additional one or two amino acids inserted are hollow but with three or more amino acids show a protein density in the center. Black arrowheads mark the inner diameter and white arrowheads mark the outer diameter. Scale bar is 50 nm. **b** Plot of summed intensities of the micrographs in **a**. The inner diameter is marked with black arrowheads and the outer diameter with grey arrowheads. **c** Immunoblots against Hcp and VipB of the samples in **a**.

To identify the proteins that were associated with the mutant sheaths, we purified the VipA-N1,2,3,5 and 5(GA). Proteins in the purified sample were identified using mass spectrometry analysis (Supplementary Table 1). Besides VipA and VipB proteins, VipA-N3, VipA-N5 and VipA-N5(GA) sheath preparations contained large amounts of Hcp, as further confirmed by western-blot (Fig. 3c, lower panel). The presence of Hcp in these mutant sheath samples explains the solid appearance on negative stain EM (Fig. 3a). Interestingly, components of T6SS baseplate (TssE, TssF, TssG and TssK) as well as VgrG tip components were also identified in VipA-N3, VipA-N5 and VipA-N5(GA) sheath samples (Supplementary Table 1).

### Mutant sheaths assemble only from functional baseplate and non-contractile phenotype is dominant

To exclude the possibility that the non-contractile sheaths are aberrant polymers that assemble independently of other T6SS components, we imaged their assembly in a strain that lacks baseplate component TssE. In agreement with the previous observations that TssE is required for efficient sheath assembly [1,25], the frequency of wild-type T6SS sheath assembly decreased by 140-fold in the absence of *tssE*. Similarly, the assembly of D333A or VipA-N5 sheaths was clearly dependent on the presence of a functional baseplate since the number of structures assembled in the cells lacking *tssE* was reduced by at least 100-fold (Fig. 4). This indicates that stable mutant sheaths assemble from a baseplate similarly to the wild-type sheaths.

**Figure 4.**
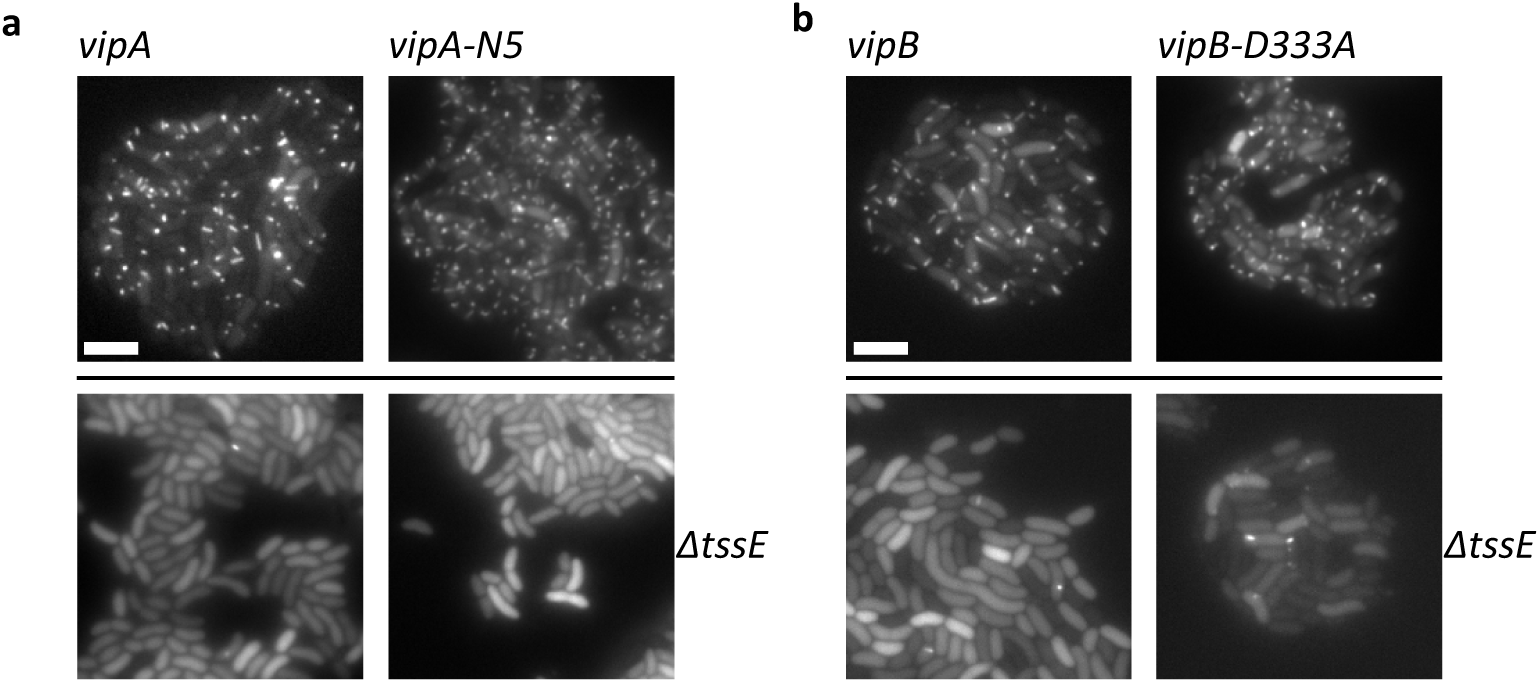
Assembly of non-contractile sheaths depends on presence of TssE. **a** Fluorescence microscopy images of V. *cholerae vipA^-^* or *vipA^-^, tssE*complemented with *vipA-msfGFP* or *vipA-5aalinker-msfGFP* on pBAD24. **b** Fluorescence microscopy images of *V. cholerae vipA^-^, vipB^-^* or *vipA^-^, vipB^-^, tssE*complemented with *vipA-msfGFP* on pBAD24 and *vipB* or *vipB-*D333A on pBAD33.

To test if the mutant sheath subunits can block T6SS activity also in the presence of the wild-type subunits, we induced expression of the VipA-N5 mutant from pBAD24 plasmid in a strain expressing wild-type sheath from the chromosome and measured efficiency of *E. coli* killing. Low level induction of VipA-N5 by 0.01% arabinose decreased T6SS-dependent killing of *E. coli* by 100- fold. The T6SS activity was almost completely blocked by increasing the concentration of arabinose to 0.1% (Fig. 5) indicating that the ratio of wild-type VipA to VipA-N5 is important for effector delivery. No such inhibition was observed when wild-type VipA was expressed from the plasmid (Fig. 5). Similarly, low level expression of VipB-D333A decreased T6SS activity by 100-fold and high level of expression blocked the T6SS activity completely (Fig. 5). This dominant negative phenotype suggests that the mutant subunits are structurally compatible with the wild-type subunits, co-assemble into the same structures and thus block T6SS function.

**Figure 5.**
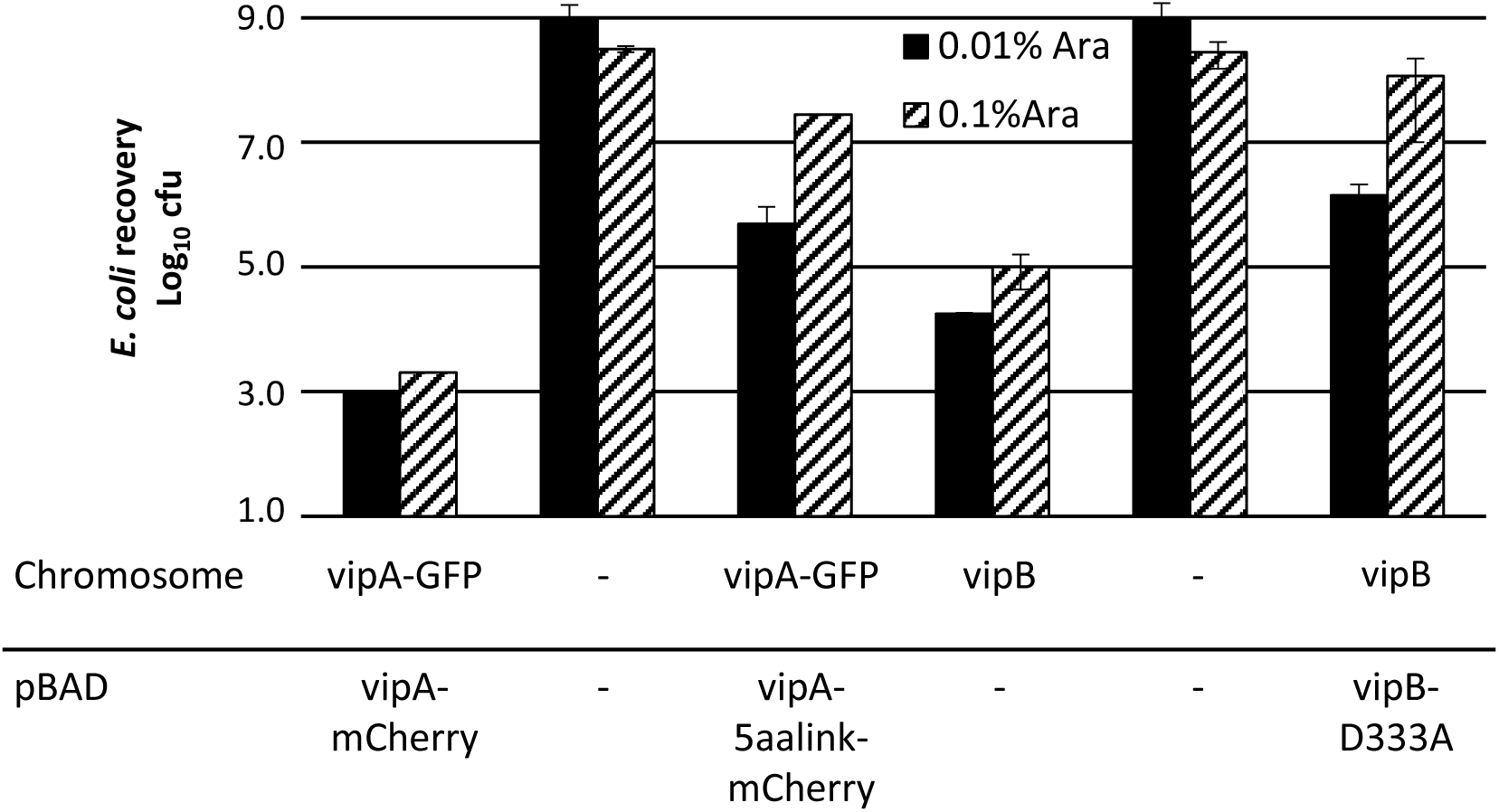
Sheath mutations show dominant negative phenotype. *E. coli* survival (± SD, N=2) after 3h competition with indicated *V. cholerae* strains in a 1:10 ratio on plate at two different arabinose concentrations.

## Discussion

T6SS is a highly dynamic system and this complicates detailed biochemical and biophysical characterization of its mode of action. Here we show that the system can be locked in the pre-contraction state by mutagenesis of the linker connecting sheath subunits or changing interactions contributing to the stability of the contracted state. Importantly, some non-contractile sheaths are stable during isolation from cells, assemble around Hcp tube, associate with many T6SS baseplate components, and co-assemble with wild-type extended sheath. This suggests that the structure of non-contractile sheaths is very similar, if not identical, to the wild-type sheath. Moreover, similarly to the wild-type extended sheath, the non-contractile sheaths form in the presence of ClpV and are therefore different from the previously described polysheath-like structures, which form independently of other T6SS components but only in the absence of ClpV [8].

Early electron micrographs of partially contracted T4 phage particles suggested that sheath contraction progresses in a wave of contracting sheath rings from the baseplate towards the phage head [24]. However, it is currently unclear how contraction of one sheath ring triggers contraction of the next ring. As we show here, insertion of two residues into the VipA N-terminal linker prevents T6SS sheath contraction *in-vivo*, suggesting that the exact length of the linker connecting the subunits is essential for sheath contraction initiation or propagation of the contraction along the sheath (Fig. 6 and Supplementary Fig. 1). T6SS, T4 and R-type pyocin sheath structures identified charged residues important for stability of contracted structures [18,21,22]. Many of these interactions specifically form during contraction and were thus proposed to be driving sheath contraction [18,21]. Together with our data, this suggests that the energy gained by formation of new charge interactions is potentially used to pull on the linkers between sheath subunits and thus propagating the contraction to the next sheath ring.

**Figure 6.**
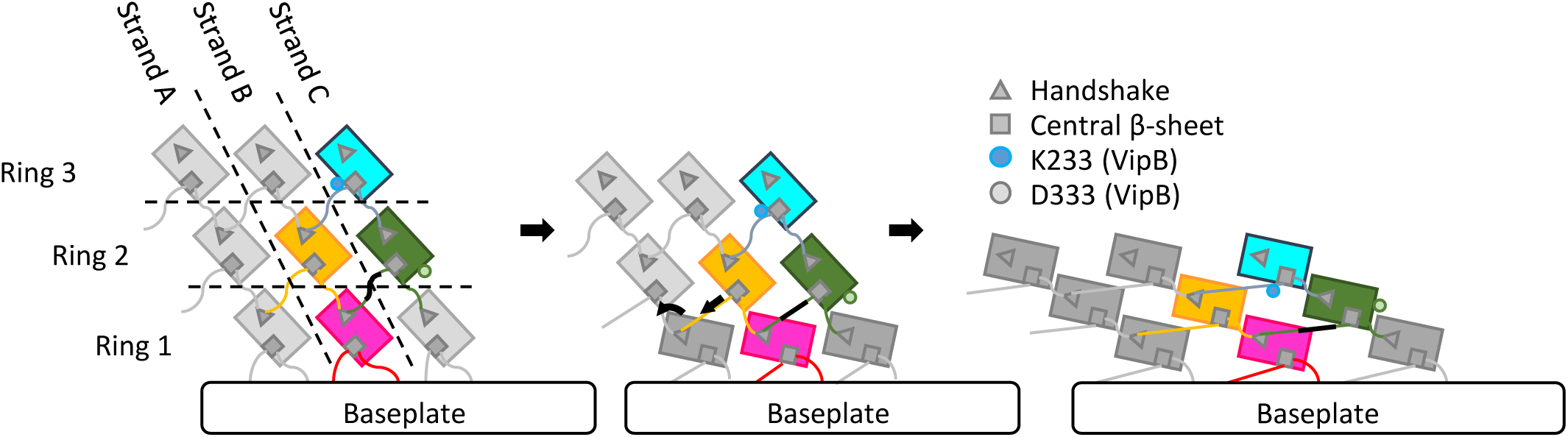
Model of the mechanism of contraction of contractile sheaths. Scheme representing the connections between strands and rings in pyocin and T6SS sheaths. Ring 1 contracts after an initial trigger coming from the baseplate via TssE. Ring 1 pulls on a linker region (black) of ring 2 and by this propagates the contraction throughout the sheath.

Interestingly, the stable non-contractile sheaths also associated with many baseplate components unlike contracted sheaths (Supplementary Table S1) [1,22]. This suggests that after sheath contraction the baseplate is destabilized and dissociates from the sheath. Live-cell imaging of sheath dynamics and localization indeed showed that few seconds after contraction, but before disassembly, sheaths often dissociate from the initial cell envelope attachment site [15]. This is consistent with the observation that T6SS assemble repeatedly inside cells and the components of the baseplate are likely reused for new rounds of assembly. In related contractile nanomachines, which are only used once, the contracted sheaths remain stably associated with the baseplates and likely provide mechanical stability to the contracted particles. In the case of contractile phages, the sheath connects baseplate and the phage head as the DNA is translocated and in the case of R-type pyocin, the sheath might be needed to stabilize the tube, which allows ion leakage and the killing of target bacterial cell [9,21,26].

The approach used here to stabilize the pre-contraction state of T6SS will be likely invaluable for further attempts to dissect T6SS mode of action at the molecular level in various bacteria and may be also used to study related contractile nanomachines with a major relevance for viral infection, bacterial competition and pathogenicity.

## Material Methods

### Bacterial strains and DNA manipulations

*V. cholerae* 2740-80 parental, *vipA-msfGFP*, *ΔvipA, ΔvipB, ΔtssE* strains and the pBAD24-*vipA-sfGFP* plasmid were described previously [1,22]. *vipA* mutants on pBAD24 plasmid were generated using standard techniques. Mutant *vipA* genes encode “A, AE, AEV, AEVE, AEVEL, AGAGA, AEVELP or AEVELPL” residues inserted right after residue 25 of wild-type *vipA*. The insertions represent either duplication of the native sequence or a sequence encoding “AGAGA”. *V. cholerae* 2740-80 *ΔvipA*-*vipB* strain was created by replacing *vipA* and *vipB* with a gene encoding “MSKEGSVGRLDQA” peptide (first seven residues of *vipA* and last six residues of *vipB* fused in frame) and *V. cholerae* 2740-80 *ΔvipA-vipB-tssE* strain was created by replacing *vipA*, *vipB* and *tssE* with a gene encoding “MSKEGSVRKYRVF” peptide (first seven residues of *vipA* and last six residues of *tssE* fused in frame) by allelic exchange as was done previously. *vipB* (wt) was cloned into pBAD24 and pBAD33 plasmids using standard techniques. K223A and D333A mutations were introduced into *vipB* using mutagenic primers. All PCR-generated products were verified by sequencing. Plasmids were transformed into *V. cholerae* by electroporation.

Gentamicin-resistant *E. coli* MG1655 strain with pUC19 plasmid was used in bacterial killing assays. Antibiotic concentrations used were streptomycin (50 μg/ml), ampicillin (200 μg/ml), chloramphenicol (20 μg/ml) and gentamicin (15 μg/ml). Lysogeny broth (LB) was used for all growth conditions. Liquid cultures were grown aerobically at 37°C.

### Bacterial killing assay

*V. cholerae* 2740-80 strains as indicated and *E. coli* MG1655 with empty pUC19 plasmid were incubated overnight at 37°C in LB supplemented with appropriate antibiotics. Cultures were diluted 100-fold, and grown to OD 0.8–1.2 in presence of appropriate antibiotics and 0.01 % arabinose for strains with pBAD plasmids. Cells were washed and mixed at final OD of ≈10 in 10:1 ratio (*V. cholerae* to *E. coli*) as specified, and 5 μl of the mixture was spotted on a pre-dried LB agar plate containing 0.01 % arabinose and ampicillin or no antibiotic. After 3 h, bacterial spots were cut out and the cells were re-suspended in 0.5 ml LB. The cellular suspension was serially diluted (1:10) in LB, and 5 μl of the suspensions were spotted on selective plates (gentamicin for *E. coli* and streptomycin for *V. cholerae*). Colonies were counted after ≈16 h incubation at 30°C. Two or more biological replicates were analyzed.

### Fluorescence microscopy

Procedures similar to those described previously [15] were used to detect fluorescence signal in *V. cholerae*. Overnight cultures of *V. cholerae* carrying pBAD24 plasmid with the respective inserts were diluted 100-fold into fresh LB supplemented with ampicillin, streptomycin, and 0.01% or 0.03% arabinose and cultivated for 2.5–3.0 h to optical density (OD) at 600 nm of about 0.8–1.2. Cells from 1 ml of the culture were re-suspended in ≈50 μl LB (to OD ≈20), spotted on a thin pad of 1% agarose in LB, and covered with a glass coverslip. Cells were immediately imaged at room temperature. A previously described microscope setup was used [22]. VisiView software (Visitron Systems, Germany) was used to record images. Fiji [27] was used for all image analysis and manipulations as described previously [28]. Bleach correction was used if necessary [29]. Contrast on compared sets of images was adjusted equally. All imaging experiments were performed with three biological replicates.

### VipA/VipB sheath preparation

Overnight cultures of the indicated strains were diluted 1:1000 in 0.5 l of fresh LB supplemented with appropriate antibiotics, and then shaken at 37°C and 250 rpm to an OD of ≈ 1.2. Cells were centrifuged for 20 min at 5000 x g and 4°C, re-suspended in 20 ml PBS and centrifuged again for 30 min at 3214 x g and 4°C. Pellets were frozen until further processing. The cell pellets were thawed, re-suspended in 20 ml of TN-buffer (20 mM Tris, 150 mM NaCl, pH 8.3) and lysed by addition of (0.75x) CelLytic™ B, Lysozyme (200 μg/ml), EDTA (5 mM) and incubation at 37°C. DNase (50 μg/ml) and MgCl2 (10 mM) was added to cleave DNA. After 15 min incubation at 37°C, cell debris was removed by centrifugation for 20 min at 10,000 x g. Cleared supernatants were subjected to ultraspeed centrifugation for 1.5 h at 104,000 x g and 4°C and the resulting pellet was washed with 1 ml TN-buffer and subsequently re-suspended in 1 ml TN-buffer, insoluble material was removed by centrifugation for 1 min at 10,000 x g. The supernatant was subjected to another round of ultraspeed centrifugation for 1h at 104,000 x g and 4°C and the resulting pellet was re-suspended in 70 μl of TN-buffer for further analysis. Purity of the sample was assessed by Coomassie stained SDS-PAGE.

### Mass spectrometry

Sheath samples (30-100 μg) were reduced with 5 mM tris(2-chlorethyl)phosphate, shaking for 1h at 37°C and alkylated with 10 mM iodoacetamide, shaking for 30 min at 25°C in the dark. Proteins were digested using sequencing-grade modified trypsin (1/250, w/w; Promega, USA) overnight at 37°C. After digestion, the samples were supplemented with TFA to a final concentration of 1%. Peptides were desalted on C18 reversed phase spin columns according to the manufacturer’s instructions (Microspin, Harvard Apparatus), dried under vacuum and re-suspended in LC-MS buffer (0.15% formic acid, 2% acetonitrile in HPLC water) at ≈0.5 mg/ml. 1 μg of peptides of each sample were subjected to LC-MS analysis using a dual pressure LTQ-Orbitrap Elite mass spectrometer connected to an electrospray ion source (both Thermo Fisher Scientific) as described recently [30] with a few modifications. In brief, peptide separation was carried out using an EASY nLC-1000 system (Thermo Fisher Scientific) equipped with a RP-HPLC column (75μm × 30cm) packed in-house with C18 resin (ReproSil-Pur C18–AQ, 1.9 μm resin; Dr. Maisch GmbH, Ammerbuch-Entringen, Germany) using a linear gradient from 95% solvent A (0.15% formic acid, 2% acetonitrile) and 5% solvent B (98% acetonitrile, 0.15% formic acid) to 28% solvent B over 75 min at a flow rate of 0.2 μl/min. The data acquisition mode was set to obtain one high resolution MS scan in the FT part of the mass spectrometer at a resolution of 240000 full width at half maximum (at m/z 400) followed by MS/MS scans in the linear ion trap of the 20 most intense ions. The charged state screening modus was enabled to exclude unassigned and singly charged ions and the dynamic exclusion duration was set to 20 s. The ion accumulation time was set to 300 ms (MS) and 50 ms (MS/MS). The collision energy was set to 35%, and one microscan was acquired for each spectrum. For all LC-MS measurements, singly charged ions and ions with unassigned charge state were excluded from triggering MS2 events. The generated raw files were imported into the Progenesis LC-MS software (Nonlinear Dynamics, Version 4.0) and analyzed using the default settings. MS/MS-data were exported directly from Progenesis LC-MS in mgf format and searched against a decoy database of the forward and reverse sequences of the predicted proteome from *Vibrio cholerae* (Uniprot, Organism ID: 243277, download date: 07/11/2016, total of 3784 entries) using MASCOT. The search criteria were set as following: full tryptic specificity was required (cleavage after lysine or arginine residues); 3 missed cleavages were allowed; carbamidomethylation (C), was set as fixed modification; oxidation (M) as variable modification. The mass tolerance was set to 10 ppm for precursor ions and 0.6 Da for fragment ions. Results from the database search were imported into Progenesis and the protein false discovery rate (FDR) was set to 1% using the number of reverse hits in the dataset. The final protein lists containing the summed peak areas of all identified peptides for each protein were exported from Progenesis LC-MS and further statistically analyzed using an in-house developed R script (SafeQuant) [30]. For relative quantification of protein abundances between sheath samples, MS1 peak intensities of the query protein (Hcp, VC_A1415; VgrG3, VC_A0123; TssE, VC_A0109; TssF, VC_A0110; TssG, VC_A0111; TssK, VC_A0114) were normalized by division with the MS1 peak intensities of VipA (VC_A0107) and VipB (VC_A0108) and again normalized to the wild-type sample. Two biological replicates were analyzed.

### Negative stain electron microscopy

300-mesh copper grids were glow-discharged for 20 s, samples (5μl, protein concentration aprox. 0.1 μg/ml) were adsorbed for 1 min and blotted using Whatman #1 filter paper. The grids were washed five times with H2O, and once using 2 % uranyl acetate, followed by a 20-s staining with 2% uranyl acetate. Grids were imaged on a CM-100 microscope (Philips N.V., Amsterdam, Netherlands) equipped with a Veleta 2k × 2k camera (Olympus K.K., Tokio, Japan) at 80 kV and a magnification of 64,000×. The pixel size was 7.4 Å. Fiji [27] was used for all image analysis.

### Immunoblot analysis

5-10 μl of purified sheath samples were mixed with 1.2-2.4 μl 4x NuPAGE^®^ LDS Sample Buffer (Life Technologies). Samples were incubated for 10 min at 95°C, centrifuged, cooled and 2 μl 1 M DTT was added. Samples were heated again for 10 min at 72°C, centrifuged, loaded on 10% polyacrylamide gels and transferred to nitrocellulose membrane (Amersham Biosciences, UK). Membrane was blocked with 5% milk in Tris buffered saline (pH 7.4) containing Tween 0.1% (TBST), incubated with primary peptide antibody against Hcp (“QSGQPSGQRVHKPF”, Genscript, USA [1]), or peptide antibody against VipB, (“QENPPADVRSRRPL”, Genscript, USA [22]) for 16 hr at 4°C or 1 hr at room temperature, washed with TBST, incubated for 1 hr with horseradish peroxidase-labeled anti-rabbit antibody (Jackson ImmunoResearch Inc., USA), and washed with the recommended buffer, and peroxidase was detected by LumiGLO^®^ Chemiluminescent Substrate (KPL, Inc., Gaithersburg, Maryland, USA). Nitrocellulose membrane was stripped using Restore^TM^ Western Blot Stripping Buffer (Thermo Scientific, USA) and reprobed using the same protocol.

### Molecular analysis

Structures of PA0622 in the contracted state (PDB ID: 3J9R) [21] and of VipA and VipB in the contracted state (PDB ID: 3J9G) [22] were aligned based on their 3D-structure using UCSF Chimera [31].

## Author contributions

M.Br. generated and characterized the mutant sheaths. Isolated and purified the sheaths, performed negative stain electron microscopy, mass-spectrometry and western-blot analysis of their structure and composition. J.W. contributed to electron microscopy data analysis. M.Ba. conceived the project and analyzed the data. M.Br., J.W. and M.Ba. wrote the manuscript. All authors read the manuscript.

## Acknowledgments

The work was supported by Swiss National Science Foundation (SNSF) grant 31003A_159525 and the University of Basel. We acknowledge the Biozentrum proteomics core facility for mass spectrometry measurements.

## Supplementary Video legends

**Supplementary Video S1. VipB residue D333 is crucial for sheath contraction.**

Time lapse videos of *V. cholerae vipA-msfGFP vipB^-^*, with pBAD24 plasmid encoding VipB or VipB-D333A mutant. Phase contrast and GFP-channel are merged in the left panel and only the GFP-signal is shown in the right panel. One field of view is 19.5 x 19.5 μm.

**Supplementary Video S2. VipB residue D333 is crucial for sheath contraction.**

Long time lapse videos of *V. cholerae vipA-msfGFP vipB^-^*, with pBAD24 plasmid encoding VipB D333A mutant. Phase contrast and GFP-channel are merged. One field of view is 19.5 x 19.5 μm.

**Supplementary Video S3. Length of N-terminal linker of VipA controls sheath contraction.** Time lapse videos of *V. cholerae vipA^-^*, with pBAD24 plasmid encoding indicated VipA-msfGFP linker mutants. Phase contrast and GFP-channel are merged in the left panel and only the GFP-signal is shown in the right panel. One field of view is 19.5 x 19.5 μm.

**Supplementary Video S4. Length of N-terminal linker of VipA controls sheath contraction.** Long time lapse videos of *V. cholerae vipA^-^*, with pBAD24 plasmid encoding VipA-N5-msfGFP mutant. Phase contrast and GFP-channel are merged. Arrows mark T6SS sheaths that break and subsequently get disassembled. One field of view is 19.5 x 19.5 μm.

**Supplementary Figure 1.**
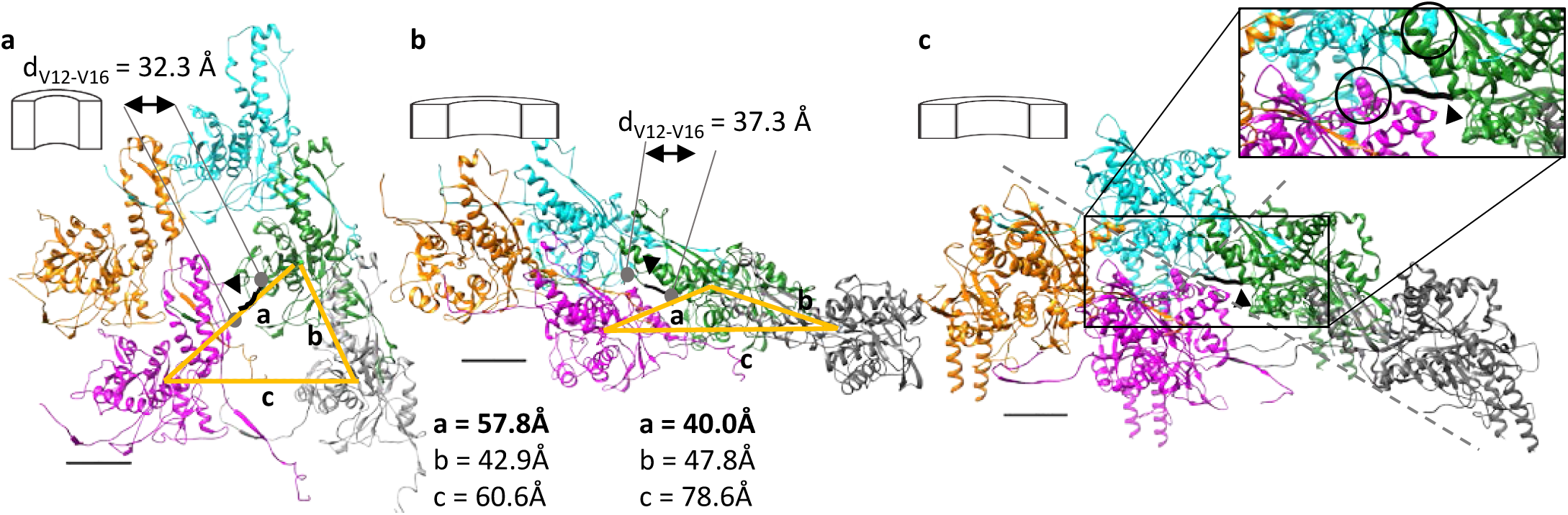
Protomers of contractile sheaths are interconnected by linkers that are stretched in the contracted state. **a** Structure of an R-type pyocin sheath in the extended state (PDB ID: 3J9Q) viewed from inside the sheath. Individual protomers are interlaced via linkers (black, arrowhead) through interactions formed by β-sheets. **b** Structure of an R-type pyocin sheath in the contracted state (PDB ID: 3J9R). The linkers interconnecting (black, arrowhead) two strands are stretched. **c** Structure of the contracted T6SS-sheath from *V. cholerae* (PDB ID: 3J9G). The backbone of the linker region of VipA that connects the green protomer with the magenta protomer is shown thick in black (arrowhead). Mutated residues on VipB are highlighted in black circles in the inset. D333 is shown on the cyan protomer, K223 is shown on the magenta protomer Distances between center of masses of different protomers are depicted as orange lines. The scale bars are 20Å.

**Supplementary Table 1.** Hcp and Baseplate Components Are Detected by Mass Spectrometry. Fold change in amounts of indicated T6SS-proteins, normalized to wild-type. Two biological replicates are shown.

|  | N1 |  | N2 |  | N3 |  | N5 |  | N5 (GA) |  |
| --- | --- | --- | --- | --- | --- | --- | --- | --- | --- | --- |
|  | Replicate 1 | Replicate 2 | Replicate 1 | Replicate 2 | Replicate 1 | Replicate 2 | Replicate 1 | Replicate 2 | Replicate 1 | Replicate 2 |
| Hcp | 0.87 | 0.94 | 1.84 | 1.53 | 12.06 | 14.54 | 29.19 | 24.07 | 19.45 | 19.74 |
| TssK | 0.97 | 0.92 | 1.46 | 0.97 | 13.45 | 4.48 | 9.64 | 3.19 | 5.13 | 5.65 |
| TssG | 0.46 | 0.65 | 0.78 | 0.71 | 21.91 | 11.36 | 9.03 | 4.79 | 6.08 | 3.93 |
| TssF | 0.71 | 0.86 | 1.32 | 0.66 | 5.04 | 4.52 | 4.95 | 2.88 | 3.44 | 3.08 |
| TssE | 0.86 | 0.82 | 0.62 | 0.76 | 5.08 | 2.94 | 4.69 | 4.79 | 2.04 | 6.78 |
| VgrG | 0.85 | 1.23 | 1.33 | 0.97 | 3.35 | 2.53 | 4.10 | 3.39 | 3.15 | 4.82 |

